# Carcinoma associated mesenchymal stem cells promote ovarian cancer metastasis by increasing tumor heterogeneity through direct mitochondrial transfer

**DOI:** 10.1101/2022.09.21.506345

**Authors:** Catherine Pressimone, Leonard Frisbie, Emma Dyer, Roja Baruwal, Claudette St. Croix, Simon Watkins, Michael Calderone, Grace Gorecki, Zaineb Javed, Huda I Atiya, Nadine Hempel, Alexander Pearson, Lan Coffman

## Abstract

Ovarian cancer is characterized by early, diffuse metastatic spread with most women presenting with extensive abdominal metastasis at the time of diagnosis. Prior work demonstrated carcinoma-associated mesenchymal stem cells (CA-MSCs) enhance ovarian cancer metastasis through a process of direct cellular interaction and formation of heterocellular CA-MSC and tumor cell complexes. In this study, we demonstrated that CA-MSCs enhance metastasis by increasing tumor cell heterogeneity through mitochondrial donation. We showed that CA-MSCs directly interacted with ovarian cancer cells via tunneling nanotubules (TNTs), and CA-MSCs used these TNTs to transfer live mitochondria to adjacent ovarian cancer cells. This mitochondrial donation preferentially occurred with ovarian cancer cells that had the lowest mitochondrial mass, as quantified using live, actively respiring mitochondrial labeling. These ‘mito poor’ cancer cells demonstrated decreased proliferation, increased sensitivity to chemotherapy, and decreased oxidative phosphorylation compared to ‘mito rich’ cancer cells. CA-MSCs rescued the phenotypes of mito poor cancer cells, restoring their proliferative capacity, increasing chemotherapy resistance, and increasing oxidative phosphorylation. We validated these findings in a fully autologous system using CA-MSCs and cancer cells derived from the same patient to prevent confounding effects of cellular response to foreign organelle/DNA. Using a knockdown of the mitochondrial motor protein, MIRO1, in CA-MSCs, we demonstrated that mitochondrial transfer is necessary for the CA-MSC-mediated rescue of ‘mito poor’ cancer cells. Mitochondria of CA-MSC origin persisted in tumor cells over multiple passages. Importantly, CA-MSC mitochondrial donation occurred *in vivo*, significantly enhanced tumor cell heterogeneity and decreased survival in an orthotopic ovarian cancer mouse model. Collectively, this work identified CA-MSC mitochondrial transfer as a critical mediator of ovarian cancer cell survival, heterogeneity, and metastasis, and blocking CA-MSC mitochondrial transfer represents a unique therapeutic target in ovarian cancer.

## Introduction

Ovarian cancer is the deadliest gynecologic malignancy in the Western world, notorious for its difficult detection, rapid metastatic spread, and development of chemotherapy resistance^1^. A major mediator of metastasis and chemotherapy resistance is genetic heterogeneity of cancer cells within the tumor^2,3,8^. Enhanced tumor cell clonal heterogeneity has been shown to increase cancer cell survival and is an independent prognostic factor for decreased survival in ovarian cancer patients^9,10,11^. The mechanisms which enhance tumor cell clonal heterogeneity in ovarian cancer are unclear, though increasing evidence suggests that the tumor microenvironment plays a critical role in shaping tumor cell behavior at both the primary tumor and metastatic sites.

Our previous work identified carcinoma-associated mesenchymal stromal/stem cells (CA-MSCs) as an essential component of the ovarian tumor microenvironment supporting ovarian cancer growth, survival, chemotherapy resistance, and metastasis ^4,5,13,15^. CA-MSCs are multipotent stromal cells that originate from epigenetic reprogramming of resident tissue MSCs, inducing a phenotypic change which conveys pro-tumorigenic properties^4,6,15^. A critical consequence of this pro-tumorigenic education is the high affinity of CA-MSCs to bind to adjacent ovarian cancer cells^4^. We have shown that CA-MSCs directly bind to ovarian cancer cells forming heterocellular complexes which co-metastasize to distant organs^4^. The physical interaction between CA-MSCs and ovarian cancer cells appears to be a critical mediator of metastasis, though the precise mechanism remains undefined.

MSCs are known to interact with adjacent cells in many ways including donation of mitochondria in times of stress. MSCs have been shown to donate mitochondria and rescue cardiomyocytes^38, 39^ and neurons that are experiencing hypoxia^20-23^. Mitochondrial transfer for metabolic rescue has also been demonstrated in cancer with immune cells donating mitochondria to breast cancer cells.^6^ This mitochondrial transfer occurs via physical interaction between the cancer and immune cells, mirroring the heterocellular complexes that form between CA-MSCs and ovarian cancer cells. The ovarian tumor microenvironment is notoriously hypoxic and depleted of nutrients^24-36^. We thus hypothesized that CA-MSCs donate mitochondria to ovarian cancer cells to overcome this stress, enhancing tumor cell heterogeneity and metastasis.

Here we demonstrated that CA-MSCs donate mitochondria to ovarian cancer cells that have the least endogenous mitochondrial bulk, rescuing these ‘mito poor’ cancer cells by restoring their oxidative phosphorylation capacity, enhancing proliferation capability, and reducing chemotherapy sensitivity. Conversely, blocking mitochondrial transfer by knocking down the mitochondrial motor protein, MIRO1 (mitochondrial Rho GTPase 1), prevented the functional and metabolic rescue of tumor cells by CA-MSCs, verifying that this phenotypic rescue is mediated through mitochondrial transfer. We further demonstrated that blocking CA-MSC mitochondrial transfer significantly decreases metastasis, decreases clonal heterogeneity of ovarian cancer cells both at the primary and metastatic site, and improves survival in an orthotopic mouse model of ovarian cancer.

Collectively, this work illustrates a critical mechanism of stromal support of ovarian cancer metastasis. This study also goes beyond previous work by demonstrating that transferred mitochondria persist inside recipient tumor cells and increase tumor cell heterogeneity and metastasis.

## Results

### CA-MSCs increase ovarian cancer cell heterogeneity during metastasis

Previous work has demonstrated that CA-MSCs enhance tumorigenicity and metastasis of ovarian tumor cells (TCs) *in vitro* and *in vivo* ^4-^*7*. To elucidate if CA-MSCs enhance tumorigenicity and metastasis via enrichment of tumor cell heterogeneity, a DNA barcoding system was incorporated into the high grade serous ovarian cancer cell line OVCAR3, such that each TC contained one unique genetic barcode, enabling identification of individual TC clones via DNA sequencing (see methods). CA-MSCs were extracted from patient high grade serous ovarian cancer samples without prior treatment and validated as previously described^6^. Using a tail-vein injection model, TCs alone or TCs + CA-MSCs were injected into NSG mice. CA-MSCs were injected in a 1:10 ratio with TCs based on our prior work demonstrating equivalent CA-MSC:TC ratios in human ascites complexes^4^. A tail-vein model was used to bypass confounding effects of differential primary tumor growth, thus enabling unadulterated assessment of clones that have entered the circulation. Additionally, previous work highlighted that hematogenous metastasis creates more selective pressure for abdominal colonization, and this is more efficiently achieved with a tail-vein model when compared to an intraperitoneal model^13^. After 21 days, when the first mouse met endpoint criteria, all mice were euthanized and sites of metastasis were quantified at necropsy. Liver, lung, and blood were isolated and used for DNA sequencing to identify ovarian TC clones. While both TC alone and TC + CA-MSC groups demonstrated 100% lung metastasis, the TC + CA-MSC group demonstrated enhanced liver metastasis (100% vs 57%), abdominal metastasis (100% vs 0%), and malignant ascites (100% vs 63%) when compared to the TC-only group (Fig1A). TC + CA-MSC co-injection versus TC alone resulted in significantly higher clonal heterogeneity within metastatic sites such as the liver, lung and blood (Fig1B-D). Furthermore, as demonstrated in Figure 1E, CA-MSC co-injection with TCs resulted in a greater total number of unique TC clones within each metastatic site. When normalized for tumor content, the number of unique barcodes per microgram of tumor DNA remained higher in the metastatic sites of mice injected with TC+CA-MSCs compared to TC alone samples (Sup table 1). As shown in Figure 1F, the distribution of unique barcodes is increased across all metastatic sites in TC+CA-MSC groups compared to TC alone groups. As an additional quantification of tumor cell clonal diversity, we also compared the number of cells that arise from the top 50 most abundant clones/barcodes at each primary and metastatic site in the TC and TC+CA-MSC groups (Fig1G). The TC+CA-MSC sites demonstrated a greater number of unique clones compared to TC alone indicating that the presence of CA-MSCs improved survival and propagation of a larger pool of cancer cell clones.

**Figure 1:**
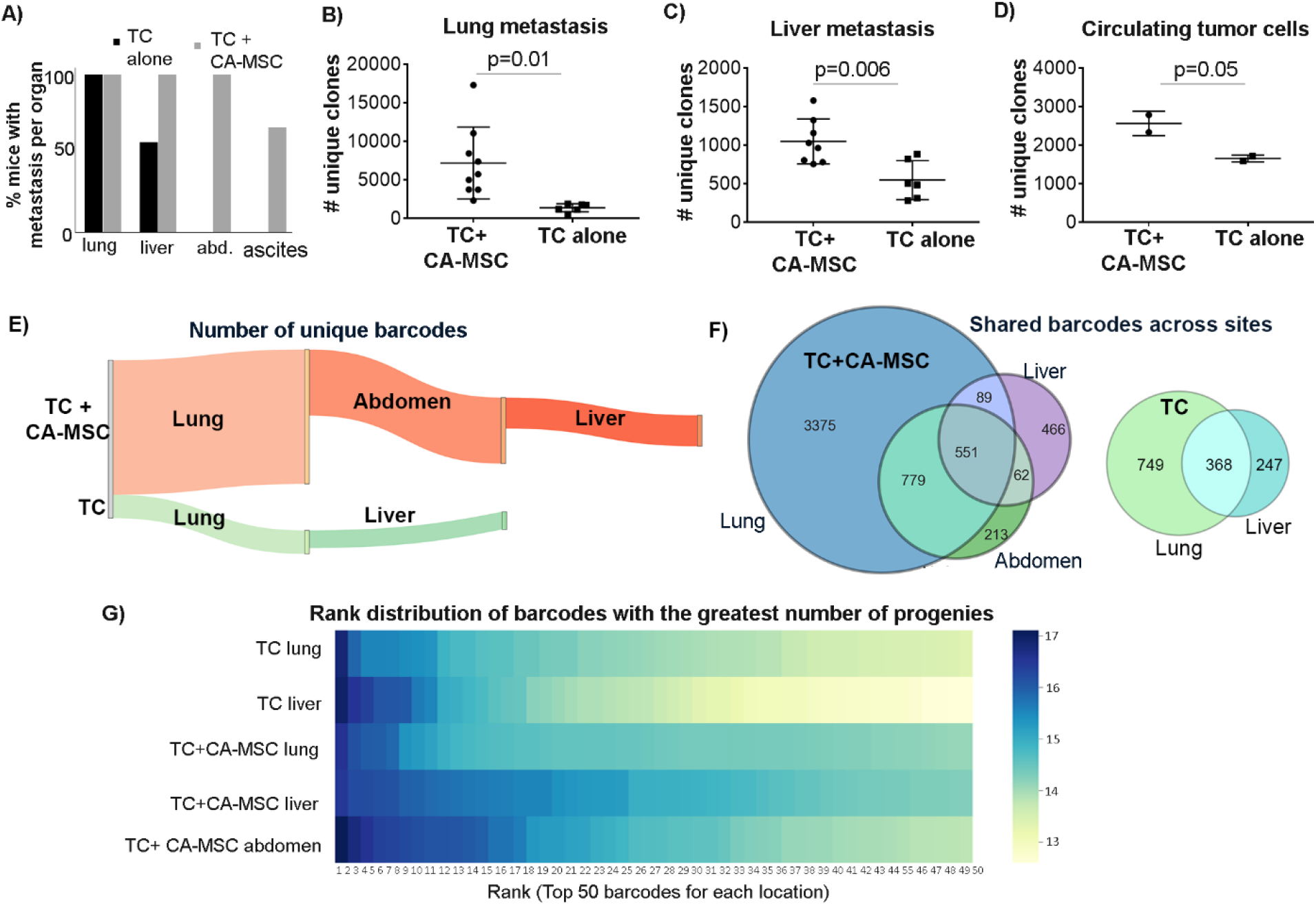
CA-MSCs increase OVCAR3 ovarian tumor cell heterogeneity at metastatic niches *in vivo*. A) Quantification of organ specific metastasis per mouse group: tumor cell (TC) alone vs TC + CA-MSC. B-D) Quantification of the number of unique clones at each site of metastasis (B-lung; C-liver; D-blood). E) Sankey diagram visually demonstrating the number of unique barcodes at each tumor site in TC+CA-MSC vs TC alone animals. F) Venn diagrams demonstrating the distribution of the tumor cell clones across metastatic sites per animal. G) Heatmap ranking the log-based total number of unique clones derived from the top 50 most abundant barcodes at each metastatic site, demonstrating that tumors containing CA-MSCs had more abundance of unique progeny and thus increased clonal heterogeneity.

### CA-MSCs transfer mitochondria to adjacent tumor cells

We demonstrated that CA-MSCs increase ovarian cancer metastasis and clonal heterogeneity, but the mechanisms driving this phenotype remain unclear. Our previous work demonstrated that direct contact between CA-MSC and TCs was necessary to enhance metastasis^4^. We therefore investigated if this direct interaction also drives tumor cell heterogeneity. We stably labeled CA-MSCs mitochondria via with lenti-viral transduction of GFP-COX8 which becomes incorporated into all mitochondria. These GFP-COX8 CA-MSCS were grown with TCs fluorescently labeled with cellTrace blue and monitored with real time IncuCyte imaging. Within 12 hours, CA-MSCs and TCs readily formed tunneling nanotubes (TNTs), suggesting that physical contact via TNTs is a significant interaction between CA-MSCs and TCs (Fig 2A). Previous studies have shown that MSCs use TNTs to donate mitochondria to adjacent cells in times of ischemic and metabolic stress ^20, 21, 22, 23, 25^, which are similar conditions to the hypoxemia and nutrient scarcity of the ovarian tumor microenvironment ^24-36^. To investigate if CA-MSCs donate mitochondria to adjacent ovarian tumor cells via physical interaction, we repeated the IncuCyte experiment along with a red actin stain to highlight TNT formation. Real-time Incucyte imaging illustrated green CA-MSC mitochondria were transferred to adjacent ovarian TCs via TNTs, solidifying that mitochondrial transfer occurs during direct CA-MSC and TC interactions.

**Figure 2:**
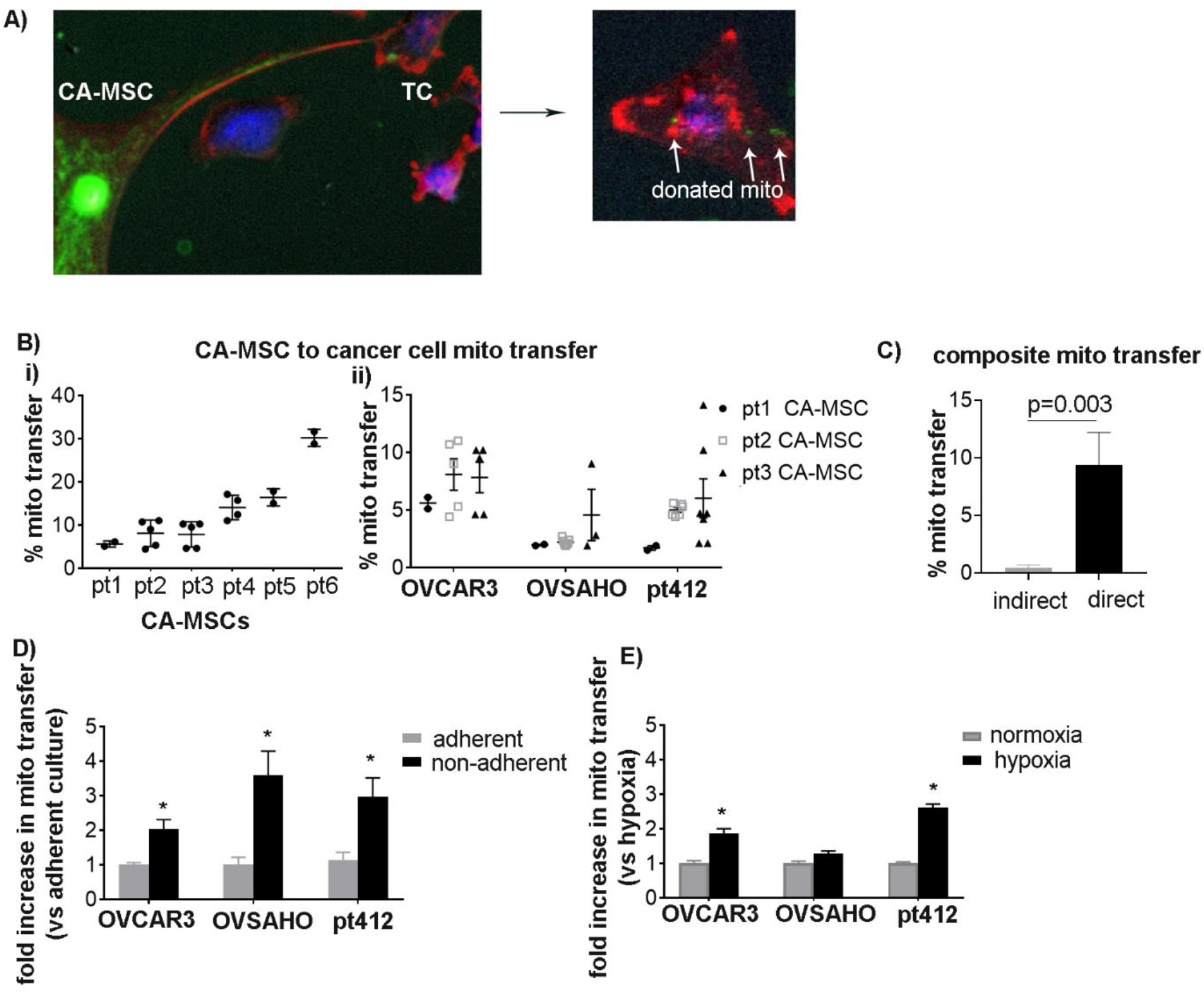
CA-MSCs directly transfer mitochondria to adjacent ovarian tumor cells. A) Imaging. B) Transfer of GFP-labeled CA-MSC mitochondria to tumor cells was quantified using flow cytometry. Six individual patient derived CA-MSCs are represented (i) and three individual patient derived CA-MSCs donating to three individual ovarian cancer cell lines are represented (ii). C) CA-MSC to tumor cell mitochondrial transfer under indirect co-culture vs direct co-culture is quantified (composite of 5 individual experiments). D) Change in CA-MSC to tumor cell mitochondrial transfer under adherent vs non-adherent conditions (composite of 5 individual experiments). E) Change in CA-MSC to tumor cell mitochondrial transfer under normoxic (21% O2) vs hypoxic (1% O2) conditions. In B-D, CA-MSCs were co-cultured in a 1:1 ratio with TCs. *=P<0.05. Mean and SEM are plotted.

To determine if the mitochondrial transfer findings are generalizable to all CA-MSC cells, we tested the ability of 6 independent, patient-derived CA-MSCs to donate mitochondria to three unique ovarian cancer cell lines: two well established high grade serous lines (OVCAR3, OVSAHO) and one primary patient-derived high grade serous line (pt412). We directly co-cultured GFP-COX8 transduced CA-MSCs with cellTrace blue-labeled cancer cells for 24 hrs. After co-culture, cells were dissociated into single cell suspension and flow cytometry was used to quantify the number of TCs which gained CA-MSC mitochondria, based on cellTrace blue and GFP positive gating. All six patient-derived CA-MSC lines were capable of donating mitochondria to each cancer cell line tested. However, the amount of mitochondrial transfer varied amongst different CA-MSC donors (Fig 2B). Mitochondrial transfer between CA-MSCs and TCs requires direct physical contact, as exhibited by the near absence of mitochondrial transfer when CA-MSCs and TCs were only allowed to indirectly interact through a transwell system or with CA-MSC conditioned media (Fig2C). The necessity of direct physical interaction for mitochondrial transfer further substantiates that tunneling nanotubules may be a main mechanism by which CA-MSCs transfer mitochondria to TCs.

Given that CA-MSCs donate mitochondria to adjacent TCs via direct interaction, we explored if other physiologic conditions that are intrinsic to the ovarian tumor microenvironment may influence mitochondrial transfer. Tumors exist in three-dimensional orientations, with cells interfacing in multiple planes. Additionally, to effectively metastasize, TCs must detach from the primary tumor and survive under non-adherent conditions. Indeed, our previous work demonstrated that CA-MSCs directly interact with TCs to form heterocellular spheroids which enhance metastasis and exist within ovarian cancer malignant ascites^4^. We expanded on that knowledge by testing if CA-MSC mitochondrial transfer was impacted by non-adherent, spheroid conditions. We found that TCs cocultured 1:1 with CA-MSCs under non-adherent conditions exhibited a two-to-four-fold increase in mitochondrial transfer compared to cocultures in standard adherent conditions (Fig2D). Given that MSCs have been shown to donate mitochondria under ischemic stress to cardiomyocytes^20, 21, 22, 23^ and that the ovarian tumor microenvironment is hypoxic^24-36^, we also tested if hypoxia influenced CA-MSC mitochondrial transfer. CA-MSCs and TCs were grown under non-adherent conditions in normoxic (21% O2) or hypoxic (1% O2) conditions. We demonstrated that 24 hours of coculture in hypoxia enhances CA-MSC mitochondrial donation to OVCAR3 and pt412 tumor cell lines two-fold compared to normoxia (Fig2E).

We next examined whether intrinsic characteristics of the tumor cells affect mitochondrial donation from CA-MSCs (pts 3, 4 and 5 were used for subsequent experiments). We tested if the endogenous mitochondrial bulk within tumor cells altered CA-MSC mitochondrial gain. TCs were stained with MitoTracker Deep Red, a dye that is taken up by actively respiring mitochondria, and stained TCs were sorted by flow cytometry into TCs with the most (top 20%) or least (bottom 20%) amount of endogenous respiring mitochondria. We observed that TCs with the least endogenous mitochondria (referred to as ‘mito poor’ TCs) received significantly more CA-MSC mitochondria compared to TCs with the most endogenous mitochondria (referred to as ‘mito rich’ TCs) (Fig3). This phenomenon was replicated and consistent across three high grade serous ovarian cancer cell lines and 3 independent patient derived CA-MSCs (Fig 3A).

**Figure 3:**
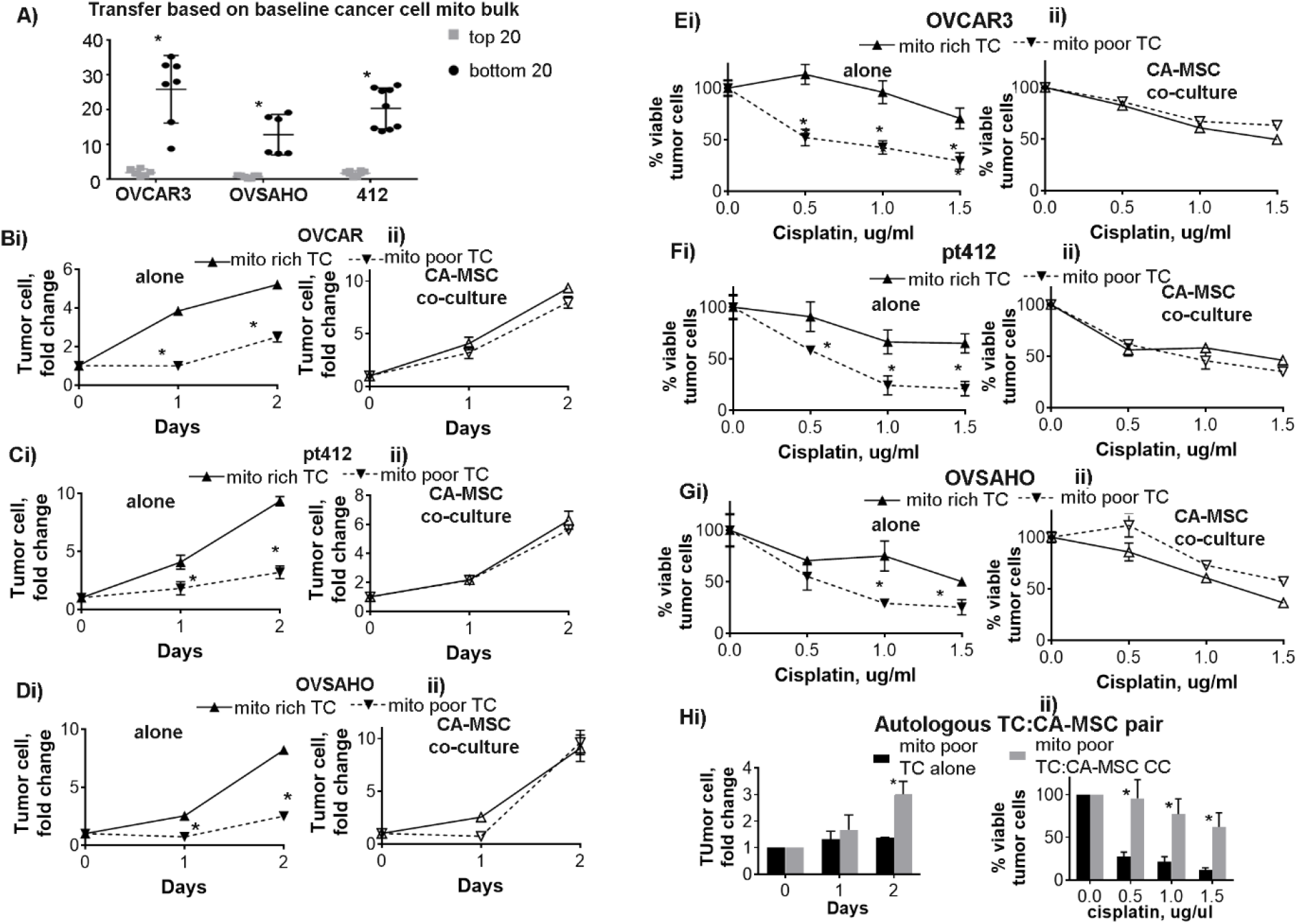
TCs with less endogenous mitochondrial bulk demonstrate decreased proliferation and increased chemotherapy sensitivity compared to TCs with high endogenous mitochondrial bulk. CA-MSC co-culture rescues ‘mito poor’ TCs, restoring proliferation and chemotherapy resistance A) CA-MSC mitochondrial transfer was quantified comparing donation to tumor cells (TCs) with the top 20% mitochondrial bulk vs bottom 20% mitochondrial bulk demonstrating significantly more transfer to the latter. B-Di) TCs with the top 20% mitochondrial bulk (‘mito rich’) vs bottom 20% mitochondrial bulk (‘mito poor’) were counted over time to assess proliferation, ii) ‘mito rich’ vs ‘mito poor’ TC were co-cultured with CA-MSCs and TC proliferation was quantified demonstrating co-culture rescues the proliferation of ‘mito poor’ TCs. Three different TC lines were used (B) OVCAR3, (C) pt412, (D) OVSAHO. *=p<0.05. Mean and SEM are represented from at least 3 independent experiments. E-G) Survival of increasing doses of cisplatin in ‘mito rich’ vs ‘mito poor’ (i) TC alone and with (ii) CA-MSC co-culture. Three different TC lines were used (C) OVCAR3, (D) pt412, (E) OVSAHO. *=p<0.05. Mean and SEM are represented from at least 3 independent experiments. H) ‘Mito poor’ TCs from a paired tumor cell and CA-MSC sample derived from the same patient with high grade serous ovarian cancer were grown alone or with CA-MSC co-culture and (i) TC proliferation and (ii) chemotherapy sensitivity were compared validating the rescue of ‘mito poor’ TC growth and chemotherapy survival with CA-MSC co-culture in an autologous paired sample. *=p<0.05. Mean and SEM are represented from 3 independent experiments.

### CA-MSC co-culture enhances ‘mito poor’ TC proliferation and chemotherapy resistance and restores their ‘metabolic fitness’

Ovarian tumor cells received mitochondria from adjacent CA-MSCs, with significantly more mitochondria being transferred to ‘mito poor’ TCs. In order to elucidate why CA-MSCs preferentially donate mitochondria to ‘mito poor’ TCs, we assessed the baseline phenotypic differences between the ‘mito poor’ TCs vs the ‘mito rich’ TCs. Again, TCs were stained with MitoTracker Deep Red and sorted into the top 20% ‘mito rich’ and bottom 20% ‘mito poor’ groups. The proliferation rate for each group was quantified through manual cell counting. The ‘mito poor’ TCs demonstrated remarkably less proliferation compared to the ‘mito rich’ TCs (Fig3B-Ei). We then assessed the proliferation of the ‘mito poor’ and ‘mito rich’ TCs following 24hrs of CA-MSC co-culture and flow cytometric sorting. The ‘mito poor’ and ‘mito rich’ TC groups were defined by their initial endogenous mitochondrial bulk not accounting for GFP-labeled mitochondria gained from the CA-MSCs. CA-MSC co-culture led to significant increases in proliferation in the ‘mito poor’ TCs, effectively restoring their proliferative capacity equal that of the ‘mito rich’ TCs (Fig3B-Eii).

Additionally, the ‘mito poor’ TCs were more sensitive to cisplatin treatment at baseline compared to the ‘mito rich’ TCs. Figure 3E-Gi demonstrates the survival of ‘mito poor’ vs ‘mito rich’ TCs to increasing doses of cisplatin. To determine the impact of CA-MSC co-culture on ‘mito poor’ vs ‘mito rich’ TCs, CA-MSCs and TCs were co-cultured for 24hrs and the ‘mito poor’ vs ‘mito rich’ TC were flow sorted as above. Sorted TCs were treated with cisplatin and viable cells counted after 48hrs. CA-MSC co-culture enhanced the chemotherapy resistance of the ‘mito poor’ TCs to a level equal to the ‘mito rich’ TCs. The proliferation and chemotherapy resistance assays for the ‘mito poor’ and ‘mito rich’ TCs with and without CA-MSC co-culture were repeated in triplicate and with three independent ovarian tumor cell lines. Collectively, these data demonstrate TCs with less endogenous mitochondria preferentially receive CA-MSC mitochondria and that co-culture with CA-MSCs enhances the proliferation and cisplatin resistance of ‘mito poor’ TCs with less endogenous mitochondria.

To ensure the mitochondrial transfer and rescue of ‘mito poor’ TCs was not an artifact of using cells from 2 different origins and thus eliciting a stress response from foreign mtDNA, we validated the above experiments using a paired patient sample. Cancer cells and CA-MSCs were isolated from the same patient with high grade serous ovarian cancer. We first verified CA-MSCs donated mitochondria to its autologous tumor cell pair. We then isolated the mito poor TC from the paired sample and tested the proliferation and chemotherapy sensitivity of the ‘mito poor’ TCs alone or after CA-MSC co-culture. Figure 3F demonstrates CA-MSC co-culture similarly enhances the proliferation and survival of autologous ‘mito poor’ TCs.

We next assessed if the mitochondria donated from CA-MSCs to TCs remain actively respiring and alter oxidative metabolism in TCs. To assess this, we measured the oxygen consumption rate (OCR) using live cell extracellular flux analysis. We first analyzed the baseline respiratory capacity of the ‘mito rich’ compared to ‘mito poor’ TCs. TCs were sorted as above and analyzed with an Agilent XF Seahorse to measure mitochondrial respiration. Figure 4A demonstrates that the ‘mito rich’ TCs had significantly higher basal respiratory capacity, maximal respiratory capacity, ATP-dependent OCR, and spare respiratory capacity compared to ‘mito poor’ TCs. However, following co-culture with CA-MSCs, the metabolic profile of the ‘mito poor’ TCs matched that of the ‘mito rich’ TCs, indicating that CA-MSCs restored the metabolic fitness of the ‘mito poor’ TCs, possibly through gain of actively respiring CA-MSC mitochondria. This data provides evidence that the CA-MSC mitochondria gained by TCs are likely metabolically active and may be enhancing the respiratory capacity of TCs that contain less endogenous mitochondria.

**Figure 4:**
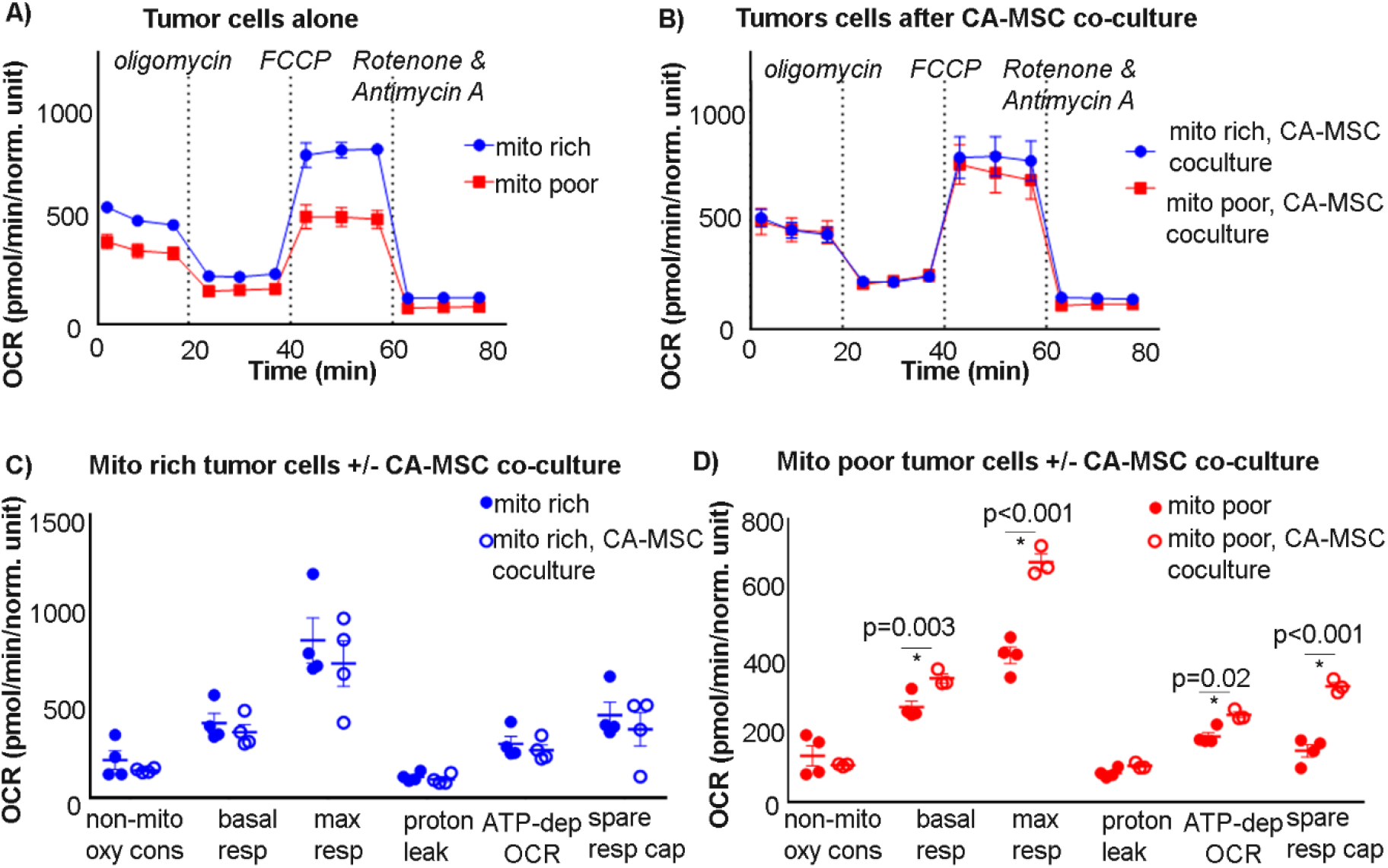
Respiratory capacity of ‘mito poor’ TCs improves after CA-MSC co-culture. A-B) Representative seahorse assay plots of ‘mito rich’ vs ‘mito poor’ tumor cells (A) alone and (B) after CA-MSC co-culture. C-D) Summary data of metabolic parameters for the ‘mito rich’ vs ‘mito poor’ tumor cells (C) alone and (D) after CA-MSC co-culture. Each point represents an independent experiment.

### MIRO1 knockdown prevents mitochondrial transfer and decreases the proliferation and chemotherapy resistance benefit of CA-MSC coculture

We demonstrated that ‘mito poor’ TCs receive CA-MSC mitochondria at higher rates and subsequently have increased proliferation, chemotherapy resistance, and oxidative respiration following co-culture with CA-MSCs. To verify this impact is due to CA-MSC mitochondrial transfer and not other confounding mediators, we targeted the mitochondrial transfer protein MIRO1. MIRO1 is a mitochondrial transport protein that facilitates intracellular movement of mitochondria via microtubules^16, 17, 18, 19^. We created MIRO1 knockdown (KD) CA-MSCs, which prevented most mitochondrial transfer while still enabling the CA-MSC to directly interact with the TC. MIRO1 was knocked down in multiple CA-MSC primary cell lines via lentiviral siRNA, with mRNA decreasing 50% and protein expression decreasing 80% (Fig5A). We validated that MIRO1 KD did not impact CA-MSC viability, growth, or ability to interact with TCs (Fig5B, Sup Fig1).

We next demonstrated that MIRO1 KD effectively prevented CA-MSC to TC mitochondrial transfer. MIRO1 KD CA-MSCs with COX8-GFP labeled mitochondria were co-cultured with fluorescently labeled TCs as above and mitochondrial transfer was quantified via flow cytometry. Figure 5C illustrates that MIRO1 KD CA-MSCs transfer 15-20 fold less mitochondria to ovarian TCs compared to CA-MSCs with wild type MIRO1 expression. We next repeated the proliferation, chemotherapy resistance, and metabolomic experiments with the ‘mito rich’ and ‘mito poor’ TCs with and without control CA-MSC or MIRO1 KD CA-MSC co-culture. MIRO1 KD CA-MSCs failed to alter the proliferation or chemotherapy resistance of the ‘mito poor’ TCs. Similarly, MIRO1 KD CA-MSCs did not alter the metabolic profile of the ‘mito poor’ TCs (Fig5F, G), indicating that mitochondrial transfer is critical for the CA-MSC mediated enhancement of ‘mito poor’ TC proliferation, chemotherapy resistance, and metabolic function.

**Figure 5:**
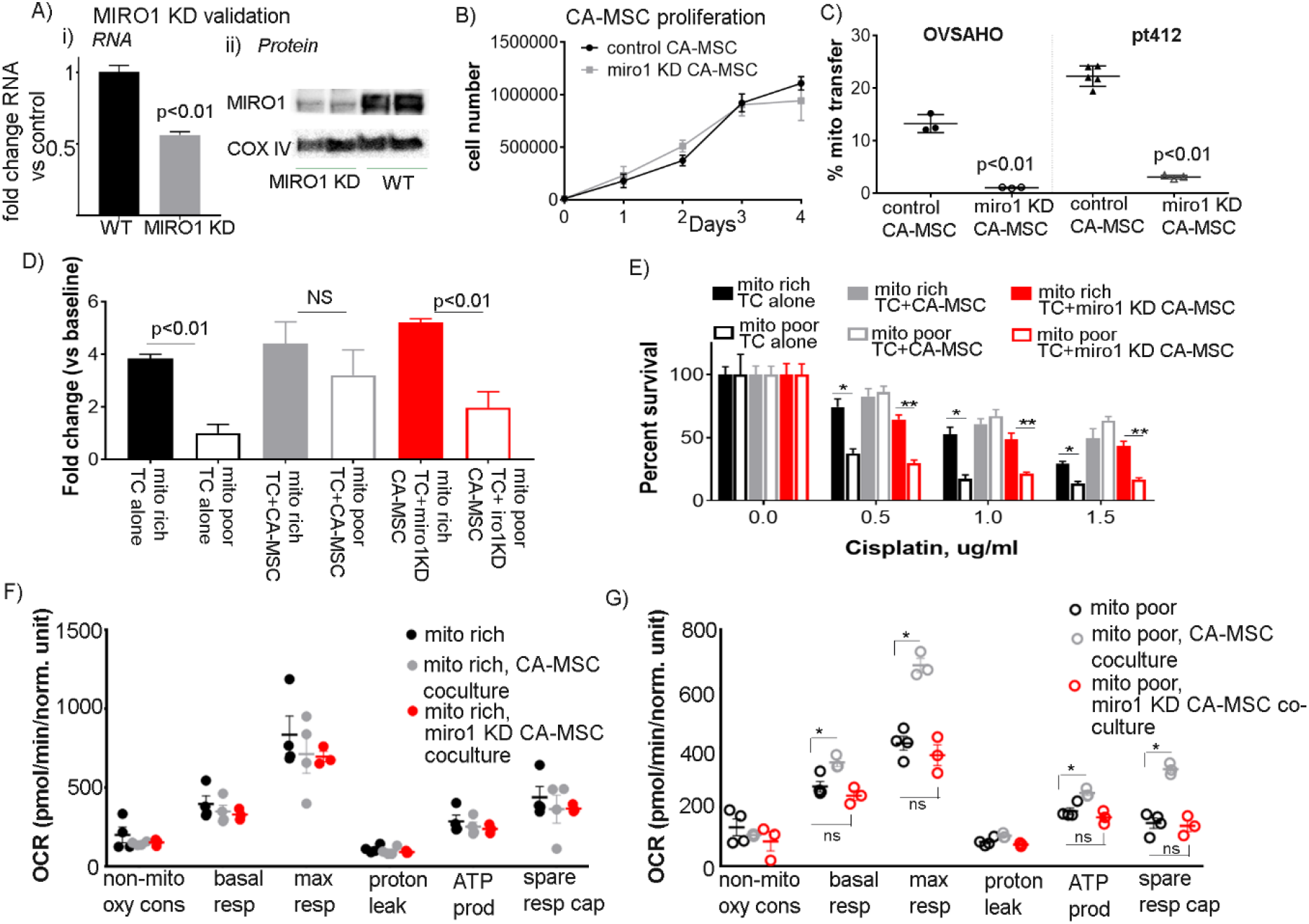
Blocking CA-MSC mitochondrial transfer via MIRO1 knockdown eliminates the proliferation and chemotherapy resistance benefit of TC coculture with CA-MSCs. A) MIRO1 KD validation at the RNA (i) and protein (ii) level in CA-MSCs. WT=wild type control. B) Proliferation of MIRO1 KD vs control CA-MSCs demonstrated that MIRO1 KD did not impact the viability or proliferation of CA-MSCs. C) Quantification of mitochondrial transfer from control vs MIRO1 KD CA-MSCs to OVSAHO or pt412 tumor cells. D) Quantification of TC proliferation, represented as fold-change from baseline at 48hrs, in ‘mito rich’ vs ‘mito poor’ TC alone and with control or MIRO1 KD CA-MSC co-culture. E) Quantification of TC survival to increasing doses of cisplatin in ‘mito rich’ vs ‘mito poor’ TC alone and with control or MIRO1 KD CA-MSC co-culture. *=difference between ‘mito rich’ vs ‘mito poor’ TC alone; **= difference between ‘mito rich’’ vs mito poor’ with MIRO1 KD CA-MSC co-culture. F) Composite metabolic data in ‘mito rich’ TC alone and with control vs MIRO1 KD CA-MSC co-culture. G) Composite metabolic data in ‘mito poor’ TC alone and with control vs MIRO1 KD CA-MSC co-culture. Each point is an independent experiment. *, **=p<0.05. Mean and SEM are represented.

**Figure 6.**
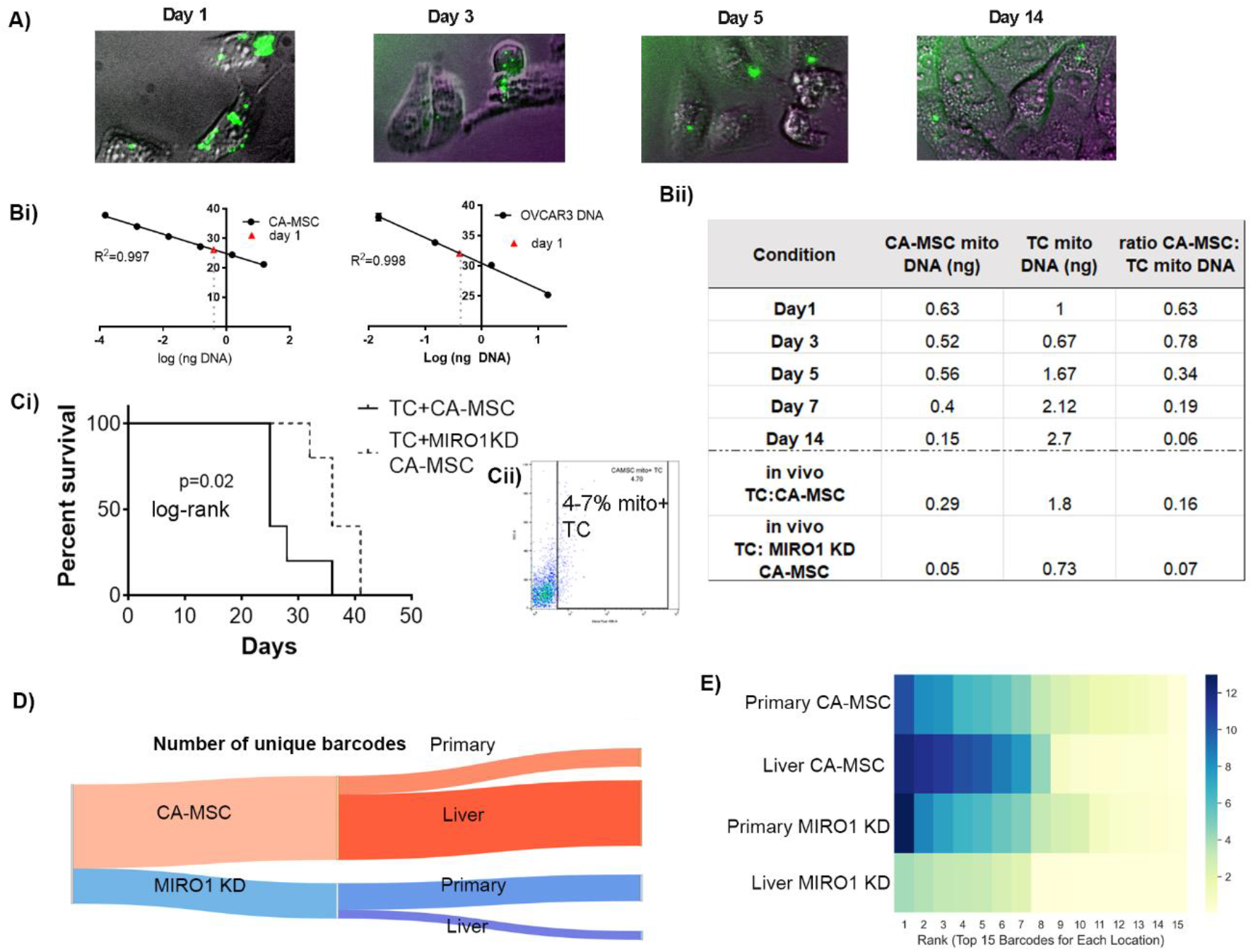
Mitochondria donated from CA-MSCs is retained after TC transfer and *in vivo* growth, and blockage of CA-MSC mitochondria transfer prolongs mouse survival and decreases TC heterogeneity in a murine ovarian cancer model. A) Immunofluorescent images of TCs which gained CA-MSC mitochondria (GFP+) after co-culture followed by flow sorting and growth of TCs alone for 1-14 days, demonstrating persistence of GFP+ mitochondria. Bi) qPCR standard curves used to quantify the amount of CA-MSC and endogenous TC mitochondrial DNA in TCs. Bii) Ratio of CA-MSC to TC mitochondrial DNA in TCs after co-culture corresponding to IF pictures in A and *in vivo* tumor cells isolated from TC:CA-MSC xenografts or TC:MIRO1 KD CA-MSC xenografts. Ci) Quantification of TC survival when TC were co-cultured with CA-MSC vs. MIRO1 KD CA-MSC. *=p<0.05. Cii) Flow cytometry analysis of isolated tumor cells with CA-MSC-donated mitochondria D) Sankey diagram demonstrating the number of unique TC barcodes in the CA-MSC vs MIRO1 KD CA-MSC xenografts. E) Heat map of the prevalence of the top 15 barcodes at each location in CA-MSC vs MIRO1 KD CA-MSC mice indicating a larger diversity of dominate clones consistent with overall increased heterogeneity in the CA-MSC groups.

We next tested to see if the donated CA-MSC mitochondria would persist or degrade after transfer to TCs. TCs that gained mitochondria from co-cultured CA-MSCs were FACS isolated and grown alone overtime. Immunofluorescent microscopy demonstrated the persistence of the transferred mitochondria over multiple days (up to 14) (Fig6A). However, this method is limited to visualizing only mitochondria that were transferred from the parent CA-MSC; the COX8-GFP tag cannot account for CA-MSC mitochondria that have replicated inside TCs as the COX8-GFP tag is nuclear encoded in the parent CA-MSC. We therefore designed haplotype-specific primers to distinguish between CA-MSC derived mitochondria (using one patient source) and TC specific mitochondria (OVCAR3, derived from a different patient) to enable the distinction and quantification of all CA-MSC derived mitochondria that existed inside TCs (originally transferred mitochondria plus transferred mitochondria that had replicated). Using quantitative PCR of the haplotype specific mtDNA, we quantified the ratio of intrinsic TC mitochondria versus donated CA-MSC mitochondria over 14 days. This method demonstrated that after 24 hours there is a 0.63 ratio of CA-MSC/TC derived mitochondrial which remains relatively stable after 3 days (0.78 ratio), decreases to 0.19 after 7 days but remains detectable at day 14 (ratio 0.06) (Fig6Bii). Given the tumor cell doubling time is ∼24hrs, this indicates not only that the CA-MSC derived mitochondria persisted inside the tumor cells, but they likely actively replicated and contributed a sizable portion of the total TC mitochondria (10-30%).

### Ovarian tumor cell proliferation, chemotherapy resistance, and clonal heterogeneity are enhanced by CA-MSC mitochondrial gain *in vivo*

We next corroborated that CA-MSC mitochondrial transfer is functionally important *in vivo* as well as *in vitro*. We used an orthotopic ovarian bursal model to more accurately model physiologic ovarian cancer development and metastasis as opposed to the earlier tail vein model which is a highly permissive metastasis model. TCs were co-injected with either control CA-MSCs or MIRO1 KD CA-MSCs into a unilateral ovarian bursa. Mice implanted with MIRO1 KD CA-MSCs demonstrated decreased metastasis and improved survival compared to mice who received control CA-MSCs (Fig6Ci). Using flow cytometry and our haplotype specific qPCR, we validated that TCs co-implanted with control CA-MSCs contained CA-MSC mitochondria at both the primary and metastatic sites. This was largely absent in the MIRO1 KD CA-MSC group (Fig6A, B).

Given the underlying hypothesis that CA-MSCs enhance metastasis through increasing TC heterogeneity by rescuing vulnerable TCs, we also employed our DNA barcoding system in the orthotopic ovarian bursa *in vivo* model. At time of necropsy, primary and metastatic tumors were removed and sent for DNA barcode sequencing. The barcoding data verified our earlier finding from the tail vein injection model that CA-MSCs enhanced the clonal heterogeneity of tumor cells compared to tumor cells grown alone (Fig6D,E). To demonstrate that the enhanced heterogeneity was due to mitochondrial transfer, the same model was used with MIRO 1 KD CA-MSCs and the tumors in MIRO1 KD mice had significantly less TC heterogeneity compared to the tumors grown in mice injected with control CA-MSCs (Fig6D, Sup Table 2).

When we quantified the cells that gave rise to the greatest number of progenies, we found that most cells in the MIRO1 KD CA-MSC tumors originated from fewer uniquely barcoded cells than in the control CA-MSC alone tumors (Fig6E). This indicates that in MIRO1 KD CA-MSC conditions, fewer unique clones were able to successfully take hold and produce a significant number of progenies as compared to control CA-MSC tumors, thereby demonstrating reduced heterogeneity in MIRO1 KD CA-MSC tumors.

## Discussion

Tumor cell viability requires functional mitochondria. The absence of mtDNA or alterations that interrupt mitochondrial proper function delay tumorigenesis^40^. Mitochondrial respiration is shown to be maintained by tumor cells in order to survive^41,42^. Metastatic cells have also been shown to rely on mitochondrial metabolism for survival^43^. Mitochondria migrate from perinuclear regions to the edges of tumor cells during metastasis which is a change that favors cell invasion by providing energy for these cells to spread into adjacent tissues ^44^. Multiple sites in this organelle have been studied as a target for cancer treatment, including structures involved in oxidative stress and electron-transportation chain^45^. Currently, a few studies involving anti-cancer therapeutics targeting mitochondrial function are in clinical Phase I trial, including in ovarian cancer patients^46^. Previous studies have shown that mitochondrial transfer from MSC to tumor cells was associated to greater oxidative phosphorylation and ATP production, resulting in tumor cell proliferation and invasion^47^. This work identified mitochondrial transfer from CA-MSCs to tumor cells as a critical mechanism mediating CA-MSC enhancement of ovarian cancer proliferation, chemotherapy resistance, tumor cell heterogeneity, and metastasis.

CA-MSCs used direct contact to donate mitochondria to ‘mito poor’ tumor cells, these cells were inherently slower growing, more sensitive to chemotherapy, and had decreased oxidative phosphorylation. CA-MSC mitochondrial donation to these cells rescued their phenotype and restored their proliferation, chemotherapy resistance, and oxidative phosphorylation capacities. Thus CA-MSC mitochondrial transfer enhanced the fitness of vulnerable ovarian cancer cells and, as a result, increased the number of tumor cell clones which survived and metastasized, resulting in increased tumor cell heterogeneity. This finding was particularly prominent at metastatic sites. While the primary tumor formed with CA-MSCs demonstrated increase tumor cell heterogeneity compared to primary tumors formed with MIRO1 KD CA-MSCs, the magnitude of difference was noticeably higher at metastatic sites (liver and abdomen). This indicated that the effect of CA-MSC mitochondrial transfer was notably important during the metastatic process and the increased fitness conveyed to the ‘mito poor’ cancer cells likely enhanced metastatic ability.

Increased cellular heterogeneity is an independent prognostic factor associated with decreased survival in ovarian cancer patients ^2,3^. Therefore, identifying a mechanism that drives tumor cell heterogeneity could offer new therapeutic targets. Indeed, diminishing CA-MSC mitochondrial transfer via MIRO1 KD resulted in improved survival in a metastatic model of ovarian cancer. In a study evaluating the role of MIRO1 in mitochondrial transfer from MSCs to injured neuron cells, MSCs expressing MIRO1 co-cultured with neurons increased cell viability and mitochondrial respiration in these affected cells. However, MIRO1 KD MSCs were not able to restore mitochondrial metabolism in neuron cells, demonstrating that MIRO1 is essential for mitochondrial donation^49^. Our finding implies that strategies blocking CA-MSC mitochondrial transfer may have important clinical implications for decreasing ovarian cancer growth, enhancing chemotherapy sensitivity, and reducing metastasis. This work expanded upon the prior reports of mitochondrial transfer. Here we quantified a subset of tumor cells which preferentially received CA-MSC mitochondria, and we demonstrated that donated mitochondria functionally impact tumor cells. Importantly, there is an inherent concern that donation of ‘foreign’ organelles or DNA can trigger a stress response which may confound the findings attributed to mitochondrial transfer. Therefore, we validated our findings in a fully autologous system using CA-MSCs and tumor cells from the same patient. Using this paired sample, we confirmed mitochondrial transfer to ‘mito poor’ tumor cells enhances proliferation and chemotherapy resistance.

Additionally, we also demonstrated that transferred mitochondria persisted in tumor cells after multiple passages and that mitochondrial transfer occurred *in vivo*. This was critical to highlight the physiologic relevance of this work. Future studies are needed to delineate the mechanisms underlying TNT formation and mitochondrial transfer in order to develop therapies that will block mitochondrial donation. Furthermore, additional work is needed to characterize the downstream consequences of CA-MSC mitochondrial transfer which extend beyond the phenotypes explored in this study.

Ultimately, this work uncovered that mitochondrial donation is a crucial mechanism of stromal support of ovarian cancer, enhancing tumor cell survival and metastasis. This both extends our understanding of critical stromal:tumor cell interactions and identifies a potentially new and powerful approach to targeting ovarian cancer metastasis.

## Materials and Methods

### In Vitro Direct and Indirect Cocultures, Normoxic and Hypoxic

CA-MSCs were grown in coculture with OVCAR, OVSAHO, or pt412 tumor cells in a one-to-one ratio. For direct adherent coculture, 5-8 × 10^5^ tumor cells and 5-8 × 10^5^ CA-MSCs were mixed in supplemented MEBM media in a T175 cm flask. For direct non-adherent coculture, 5-8 × 10^5^ tumor cells and 5-8 × 10^5^ CA-MSCs were combined and mixed with serum-free MEBM in a T75 cm nonadherent flask. For indirect coculture, 5 × 10^4^ CA-MSCs were plated on top of polystyrene 0.4um pore Transwell inserts, 5 × 10^4^ tumor cells were plated in the bottom wells, and each coculture was grown in MEBM + 10% FBS. Tumor cells grown alone at equivalent cell densities served as controls. Cocultures incubated for 24 hours in 21% O2 for normoxic experiments and 1% O2 for hypoxic experiments. Following coculture, cell types were sorted with >99% purity and used in various experiments. Non-adherent cocultures were dissociated with trypsin and mechanical agitation before sorting. To distinguish cell types for sorting, CA-MSCs were stained with cellTrace Violet dye (Invitrogen, Waltham, MA USA) in PBS suspension at 5uM for 20 minutes before neutralizing the dye with serum-containing media. CA-MSCs used in experiments were passage number 5 to 13.

### CA-MSC Mitochondria Tagging and Quantification of Mitochondrial Transfer

Early passage (<p4) CA-MSCs were transduced with a COX8-GFP lentiviral vector tagging CA-MSC mitochondria with GFP. Transduction efficiency was verified with fluorescent microscopy and flow cytometry. To identify mitochondrial transfer, tumor cells and CA-MSCs were cocultured as described above and GFP mitochondrial movement was visualized using IncuCyte real-time imaging. To quantify rates of mitochondrial transfer, tumor cells and GFP+ CA-MSCs were cocultured for 24 hours, and tumor cell gain of GFP+ CA-MSC mitochondria was assessed with flow cytometry. Cells were cocultured in adherent, nonadherent, and indirect conditions. OVCAR, OVSAHO, or pt412 tumor cell lines were each cocultured with 3 different CA-MSC lines, and the mitochondrial transfer rate of each combination was assessed in triplicate.

### Isolation of Tumor Cells with Top 20% and Bottom 20% Mitochondrial Bulk

OVCAR, OVSAHO or pt412 tumor cells were stained with MitoTracker Deep Red (Invitrogen, Waltham, MA USA), a dye of actively respiring mitochondria, in PBS suspension at 50nM for 30 minutes. Unstained tumor cells were used as controls to ensure sufficient staining occurred. Following staining, tumor cells were sorted to collect the cells with the top 20% and bottom 20% far-red fluorescence, distinguishing cells based on relative amount of endogenous mitochondrial bulk. Sorted cells were then used in various experiments.

### Proliferation and Chemotherapy Resistance Assays

OVCAR, OVSAHO or pt412 tumor cells were sorted based on endogenous mitochondrial bulk, as per above, or were sorted out of coculture with CA-MSCs, as per above, before use in the assays. Bulk tumor cells that were grown independently and sorted were used as controls. 5 × 10^3^ tumor cells were plated per well in a 96 well dish, 3-5 wells per condition, in DMEM complete media. Cells were incubated overnight to recover, and the following morning cisplatin was spiked into the media to a final concentration of 0, 0.5, 1, or 1.5uM, marking time t=0. Cells were incubated for an additional 24, 48, or 72 hours. For the experiments where tumor cells were sorted out of coculture with CA-MSCs, IncuCyte imaging was used to assess the change in cell density confluence as a proxy for cell viability. For the experiments where tumor cells were sorted based on endogenous mitochondrial bulk, cell viability was counted manually with a hemocytometer and 2:1 Trypan blue exclusion.

### Tumor Cell Sphere Formation Assay

OVCAR, OVSAHO, or pt412 tumor cells were sorted based on endogenous mitochondrial bulk, as per above, or were sorted out of coculture with CA-MSCs, as per above, before use in the assays. Bulk tumor cells that were grown independently and sorted were used as controls. 3 × 10^3^ tumor cells were plated per well in a nonadherent 96 well dish, 3-5 wells per condition, in serum-free MEBM media. Cells were incubated for 8-14 days, and the number of spheres per well was manually counted at days 5 and 8 or days 7 and 14 via compound light microscopy. Spheres were defined as suspended clusters of cells that contained >4 cells adhered to each other. (insert other materials/methods here)

Sankey diagrams, heat maps, Venn diagrams, and distribution plots were produced in Python version 3.7.3. Barcode data processing for the barcoded tail vein injection model and barcoded *in vivo* MIRO1 knockdown model was performed in Python version 3.7.3 with pandas 1.2.4 and numpy 1.20.2.

#### Sankey Diagrams

All Sankey diagrams were produced with seaborn 0.11.2 and plotly 5.7.0. In the tail vein injection model, we wanted to understand which barcodes had proliferative activity in each tumor. Therefore, we considered a barcode with reads to be any barcode with > 10 reads, indicating that at least ten cells originated from this barcode. Barcoded cells that gave rise to fewer than ten cells have minimal impact on the tumor’s heterogeneity, and the abundance of barcodes with fewer than ten reads obscure the ability to visualize the proportion of barcodes that have a greater contribution to the tumor’s heterogeneity across tumor sites. The total number of reads is the sum of all reads from all barcodes with > 10 reads.

The *in vivo* MIRO1 knockdown model Sankey diagram graphs the number of reads for each location aggregated together, after normalization to control for differences in size between tumors. This experiment used tumors from four mice, each of which had different size tumors at each site. To normalize the number of reads such that we can compare the number of reads from a smaller tumor to that of a larger tumor, the cell counts were divided by each tumor’s µg DNA. After normalizing each sample, the normalized reads were summed by condition, CA-MSC and MIRO1, as well as by site, primary and liver, to produce the total number of reads in each respective category.

#### Heat Maps and Distribution Plots

All heat map visualizations and distribution plots were produced with seaborn 0.11.2. The rank of each barcode was determined by ordering the barcodes from greatest number of progenies (1) to fewest number of progenies (*n*). The value of the upper rank *n* depends on the number of barcodes that had progenies at a particular site and condition. To account for the large distribution of counts across barcodes and visualize with a heatmap, the negative log of the cell counts associated with each barcode was calculated and plotted in rank order on the heatmap.

