## Supplemental figures and tables for "Carcinoma associated mesenchymal stem cells promote ovarian cancer metastasis by increasing tumor heterogeneity through direct mitochondrial transfer"

### Number of Unique Barcodes and Reads from Tail-Vein Injection Model

| Condition and Site | Number of Barcodes | Number of Reads | Number of Barcodes/ug DNA |
|---|---|---|---|
| TC + CA-MSC Lung | 9410 | 392918657 | 22.61 |
| TC + CA-MSC Abdomen | 4623 | 326735198 | 18.23 |
| TC + CA-MSC Liver | 2161 | 262589084 | 2.95 |
| TC Lung | 1641 | 151035028 | 5.14 |
| TC Liver | 1238 | 143062065 | 2.33 |

**Supplemental Table 1**: Shows the number of barcodes and number of total reads recorded across each site and condition in a tail vein injection model. These values correspond to the Sankey visualizations in Figure 1E-G**.**

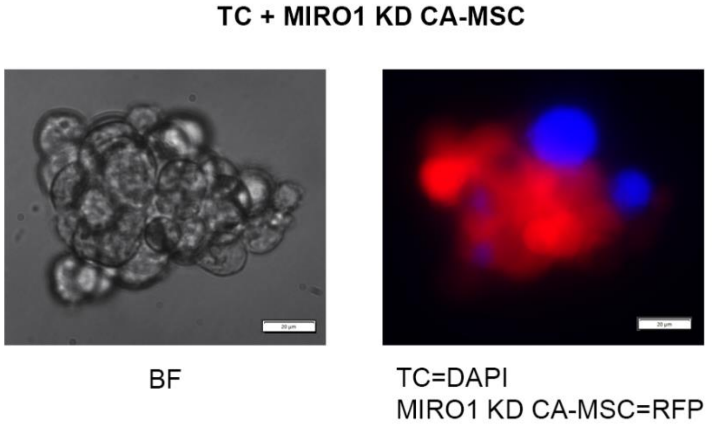


**Supplemental Figure 1. MIRO1KD CA-MSCs retain the ability to directly bind tumor cells and form heterocellular spheres under non-adherent conditions.** RFP-labeled MIRO1 KD CA-MSCs grown with cellTrace blue tumor cells directly interact forming spheroids under non-adherent conditions.

### Number of Reads in CA-MSC Alone vs MIRO1 Knockdown by Sample

| Mouse, Condition, and Site | Number of Reads | Genomic DNA in Sample (μg) | Number of Reads (normalized by amount of μg Genomic DNA by sample) |
|---|---|---|---|
| M1 CA-MSC Primary | 77886193 | 131.6 | 592065.3 |
| M1 CA-MSC Liver | 65703374 | 22.9 | 2874163.3 |
| M2 CA-MSC Primary | 73223004 | 71.4 | 1024958.1 |
| M2 CA-MSC Liver | 67607026 | 23.6 | 2870786.7 |
| M3 MIRO1 Primary | 75238821 | 42.6 | 1766169.5 |
| M3 MIRO1 Liver | 64306171 | 83.6 | 769120.6 |
| M4 MIRO1 Primary | 66617832 | 118.2 | 563745.7 |

**Supplemental Table 2**: Shows the number of reads for each site by sample in each mouse in *in vivo* MIRO1 knockdown and CA-MSC alone models. The number of reads were divided by the μg DNA in each sample to calculate a normalized number of reads such that different sites could be compared.
